# A Reproducible Dual-Model Constraint-Based Framework for Exploring Hepatic Energy Metabolism Under Stachys affinis-Derived Short-Chain Fatty Acid Scenarios

**DOI:** 10.64898/2026.03.26.714589

**Authors:** Alexander Tai Nguyen, Bryan Nguyen

**Author notes:** These authors contributed equally to this work.

## Abstract

Stachys affinis (Chinese artichoke) tubers contain 50–80% stachyose by dry weight, the most concentrated dietary source of raffinose-family oligosaccharides (RFOs) known. Because humans lack sufficient α-galactosidase activity, stachyose transits intact to the colon where microbial fermentation yields short-chain fatty acids (SCFAs). However, the quantitative impact of stachyose-derived SCFAs on host hepatic energy metabolism has not been systematically explored using genome-scale metabolic models. Three stachyose dose scenarios (Low/Mid/High: ∼25, 50, 100 g fresh tubers) were translated to SCFA availability vectors. Hepatic metabolic responses were simulated using Recon3D (10,600 reactions) and Human-GEM (13,417 reactions) under strict hepatocyte-like media, maximizing ATP maintenance flux (ATPM). FVA across multiple optimality thresholds (90–100%) and pFBA confirmed solution robustness. One-at-a-time sensitivity analysis characterized ATPM responses to individual parameter perturbations, and a ratio sensitivity sweep across six alternative SCFA profiles assessed dependence on assumed fermentation ratios. A targeted rescue experiment addressed model-specific propionate catabolism gaps. Both models showed dose-dependent ATPM increases (Recon3D: +71 to +286%; Human-GEM: +103 to +413% above baseline), with the 19–33% inter-model gap attributable entirely to Human-GEM’s functional propionate catabolism pathway. FVA confirmed near-unique optimal solutions (ATPM ranges ∼1% at 99% optimality, widening to ∼10% at 90%). Parsimonious FBA preserved identical ATPM values while reducing total flux by ∼4–14%, confirming objective robustness. SCFA ratio sensitivity across six alternative profiles showed 27– 28% ATPM variation, indicating qualitative robustness. Butyrate yielded the highest ATP per mole (∼22) in both models; propionate sensitivity was zero in Recon3D but ∼15.25 mmol ATPM/mmol propionate in Human-GEM. Reopening propionyl-CoA carboxylase (PPCOACm) in Recon3D under strict constraints converged ATPM to within 0.3–0.7% of Human-GEM, cross-validating both reconstructions. This reproducible dual-model pipeline identifies model-specific pathway gaps and provides cross-validated predictions to guide future experimental studies of how dietary SCFAs influence hepatic ATP metabolism.

## 1. Introduction

### 1.1 Stachys affinis and Raffinose-Family Oligosaccharides

Stachys affinis Bunge (Lamiaceae), commonly known as Chinese artichoke, crosne, or chorogi, is a perennial herbaceous plant cultivated primarily for its small, segmented tubers. The species has been cultivated in China and Japan for centuries, with introduction to European horticulture occurring in the late nineteenth century when tubers were brought from Beijing to the Crosne commune near Paris [1]. Although S. affinis remains a minor crop by global production volume, it has attracted increasing scientific interest due to its unusual carbohydrate profile, dominated by raffinose-family oligosaccharides (RFOs) rather than the starch that characterizes most tuber crops [1, 11].

Raffinose-family oligosaccharides are non-reducing α-galactosyl derivatives of sucrose, biosynthesized through sequential addition of galactose units from galactinol. The series begins with raffinose (trisaccharide), extends to stachyose (tetrasaccharide: Gal-Gal-Glc-Fru), and continues to verbascose (pentasaccharide). Stachyose (C24H42O21, MW 666.6 Da) is the dominant storage carbohydrate in S. affinis tubers, typically comprising 50–80% of tuber dry weight [1, 11]. This concentration substantially exceeds that of other RFO sources: soybeans contain 1–4% stachyose, lentils 2–4%, and chickpeas 1–3% [12]. This exceptional RFO density makes S. affinis the most concentrated dietary source of stachyose currently known.

The α(1,6) galactosidic linkages in stachyose render it resistant to hydrolysis by human digestive enzymes, as the human small intestine lacks sufficient α-galactosidase activity [2]. Consequently, ingested stachyose transits intact to the colon as a structurally intact oligosaccharide available for microbial fermentation, meeting the 2017 ISAPP consensus definition of a prebiotic substrate [2].

### 1.2 Microbial Fermentation and SCFA Production

Upon reaching the large intestine, stachyose enters a complex anaerobic ecosystem comprising approximately 10^11^ to 10^12^ bacterial cells per gram of luminal content [13]. Microbial α-galactosidases hydrolyze stachyose, releasing two galactose molecules and one sucrose, yielding four hexose units per stachyose molecule. These monosaccharides are fermented through central anaerobic pathways [14].

Primary degraders include Bacteroides thetaiotaomicron, which encodes α-galactosidase gene clusters within polysaccharide utilization loci [3], and Bifidobacterium species (B. adolescentis, B. longum) [15, 16]. Primary fermentation via the Embden-Meyerhof-Parnas pathway produces acetate and lactate, while propionate is generated via the succinate or acrylate pathways [4, 14]. Cross-feeding by butyrate producers (Faecalibacterium prausnitzii, Roseburia intestinalis) transforms acetate into butyrate via butyryl-CoA:acetate CoA-transferase [4, 16, 17]. Complete fermentation of one hexose yields 2–3 moles total SCFAs at ratios of approximately 60:20:20 to 75:15:10 (acetate:propionate:butyrate) [14, 18].

### 1.3 Short-Chain Fatty Acids and Host Metabolic Health

SCFAs are absorbed across the colonic epithelium via MCT1 (SLC16A1) and SMCT1 (SLC5A8), with approximately 90–95% absorption efficiency [5, 19]. Butyrate and propionate signal through G protein-coupled receptors GPR41, GPR43, and GPR109A [20, 24, 25] to modulate host appetite, insulin sensitivity, and lipid metabolism. Each SCFA has distinct metabolic fates: butyrate is the preferred colonocyte energy substrate and HDAC inhibitor [5, 22]; propionate undergoes hepatic gluconeogenesis via propionyl-CoA carboxylase and methylmalonyl-CoA mutase (B12-dependent) [5, 23]; acetate enters central carbon metabolism via acetyl-CoA synthetase for TCA cycle oxidation, lipogenesis, or ketogenesis [5, 10]. The gut-liver axis provides a direct route whereby portal-delivered SCFAs influence hepatic glucose and lipid homeostasis, contributing an estimated 5–10% of total human energy requirements [23].

### 1.4 Constraint-Based Modeling and Study Rationale

Constraint-based reconstruction and analysis (COBRA) predicts steady-state metabolic flux distributions using linear programming [6]. Flux balance analysis (FBA) identifies the flux vector optimizing a defined objective function subject to stoichiometric, thermodynamic, and capacity constraints [6, 26]. Prior human metabolic reconstructions include Homo sapiens Recon 1 [27], Recon 2 [28], and Recon 2.2 [30], each advancing network completeness and annotation quality. We employed COBRApy [7] to simulate human hepatic metabolism using two independent genome-scale reconstructions: Recon3D (10,600 reactions; 5,835 metabolites; 2,248 genes) [8] and Human-GEM (13,417 reactions; 8,378 metabolites; 3,625 genes) [31].

The use of two independent reconstructions addresses a critical limitation of single-model studies: pathway gaps and annotation differences between models can produce divergent predictions that, when identified, reveal reconstruction-specific artifacts versus genuine metabolic constraints. Where both models agree, confidence in predictions increases; where they diverge, the discrepancy illuminates specific pathway gaps that can be diagnostically investigated.

The objective function is the ATP maintenance reaction (ATPM), representing non-growth-associated maintenance energy: ATP + H2O → ADP + Pi + H+. Because hepatocytes are terminally differentiated, ATPM is more physiologically relevant than biomass maximization [6, 29]. The pipeline maps stachyose dose to SCFA vectors, imposes them as model constraints, and solves for maximal ATPM to explore how dietary S. affinis intake may influence hepatic energy metabolism under idealized steady-state assumptions. The resulting predictions are intended as quantitative hypotheses for experimental testing, not as direct forecasts of in vivo flux magnitudes.

**Figure S1.**
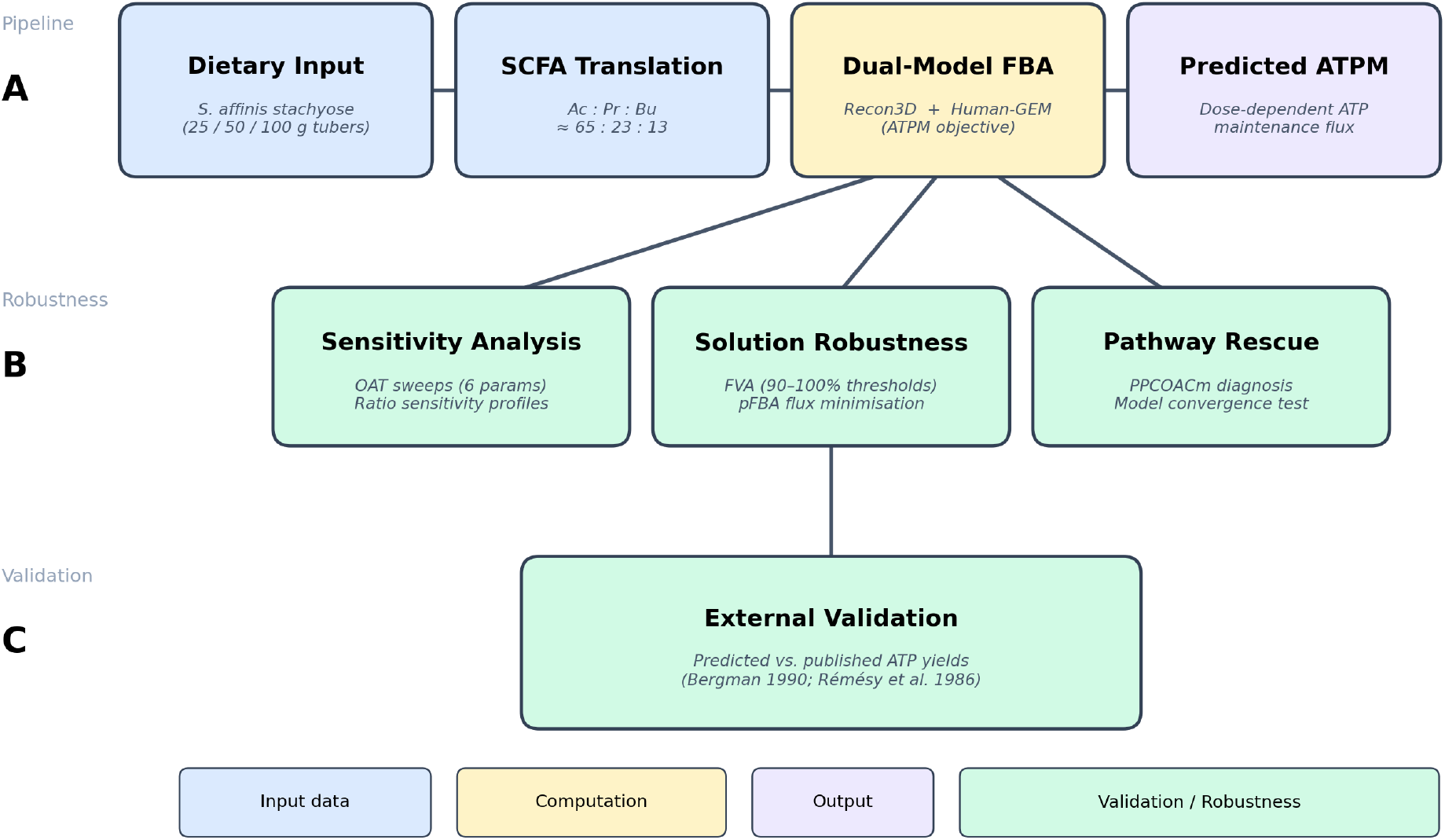
Study pipeline overview. (A) Main workflow: dietary stachyose doses are translated to SCFA availability vectors via published fermentation ratios, then simulated in two independent genome-scale models (Recon3D, Human-GEM) maximizing ATP maintenance flux. (B) Robustness analyses: one-at-a-time sensitivity sweeps, SCFA ratio sensitivity, multi-threshold FVA, and parsimonious FBA. (C) External validation: predicted ATP yields compared against published experimental measurements.

## 2. Methods

### 2.1 Study Design and Dose Scenarios

Three stachyose dose conditions were defined: StachysDose_Low, StachysDose_Mid, and StachysDose_High, corresponding to approximately 25, 50, and 100 g fresh S. affinis tubers. Assuming 20% dry matter and 60% stachyose concentration [1, 11], these correspond to approximately 3, 6, and 12 g stachyose reaching the colon. Based on published fermentation stoichiometries [14, 18], SCFA availability vectors were constructed with a fixed acetate:propionate:butyrate molar ratio of approximately 65:23:13, within the reported range for dietary fiber fermentation [18, 21]. Table 1 presents the SCFA availability vectors.

**Table 1.** SCFA availability vectors by dose condition (mmol/gDW/hr).

| Condition | Acetate | Propionate | Butyrate | Total SCFA |
| --- | --- | --- | --- | --- |
| Low | 2.0 | 0.7 | 0.4 | 3.1 |
| Mid | 4.0 | 1.4 | 0.8 | 6.2 |
| High | 8.0 | 2.8 | 1.6 | 12.4 |

### 2.2 Genome-Scale Metabolic Models

Two independent human genome-scale metabolic reconstructions were employed. Recon3D [8] encompasses 10,600 reactions, 5,835 metabolites, and 2,248 genes across eight subcellular compartments, with comprehensive coverage of central carbon metabolism, amino acid metabolism, and lipid metabolism. Human-GEM [31] is a more recent reconstruction containing 13,417 reactions, 8,378 metabolites, and 3,625 genes, with updated reaction annotations, improved pathway connectivity, and expanded coverage of cofactor-dependent metabolic routes. Both models were obtained in SBML format and loaded using COBRApy’s I/O module.

### 2.3 Simulation Protocol

An identical hepatocyte-like simulation protocol was applied to both models. The protocol comprised five steps: (1) All internal reaction bounds were capped at +/-500 mmol/gDW/hr. (2) All boundary reactions (exchange, demand, sink) were closed (bounds set to zero). (3) A curated set of essential exchange reactions was reopened: water and protons unlimited; inorganic ions at moderate bounds; O2 at 100 mmol/gDW/hr; glucose at 1.0 mmol/gDW/hr (deliberately scarce); essential amino acids and vitamins at 0.01 mmol/gDW/hr each. (4) SCFA exchange reactions were opened per the dose condition. (5) FBA was performed maximizing ATPM (capped at 500 mmol/gDW/hr).

For Human-GEM, which lacks a pre-defined ATPM reaction, an equivalent reaction (ATP_c + H2O_c → ADP_c + Pi_c + H_c) was added as ATPM_added with identical stoichiometry. Exchange reaction identifiers differ between models (e.g., EX_ac_e in Recon3D vs MAR09086 in Human-GEM); both were mapped to the same SCFA inputs. A baseline simulation with no SCFA supplementation was performed for each model.

### 2.4 Flux Variability Analysis

Flux variability analysis (FVA) was performed at 99% optimality for ATPM and key exchange reactions (SCFA uptake, O2, CO2) across all conditions in both models. FVA determines the minimum and maximum feasible flux for each reaction while maintaining at least 99% of the optimal objective value, thereby quantifying the uniqueness and robustness of the FBA solution [26]. Additional FVA at 90%, 95%, and 100% optimality thresholds is described in Section 2.10.

### 2.5 Sensitivity Analysis

One-at-a-time parameter sensitivity sweeps were performed on both models using the Mid dose as the baseline condition. Six parameters were varied independently: acetate availability (0.5– 12.0 mmol/gDW/hr), propionate availability (0.0-5.0), butyrate availability (0.0-3.0), glucose uptake (0.1-5.0), oxygen supply (10-200), and acetate:butyrate ratio (0.1-0.9, maintaining constant total carbon). For each sweep point, ATPM was re-optimized and the resulting flux recorded.

### 2.6 Propionate Pathway Rescue

To diagnose and address Recon3D’s failure to utilize propionate, a targeted pathway rescue experiment was performed. The propionyl-CoA carboxylase reaction (PPCOACm: propionyl-CoA + CO2 + ATP → methylmalonyl-CoA + ADP + Pi) was identified as blocked in Recon3D. Two rescue conditions were tested: (1) unconstrained rescue, reopening PPCOACm with default bounds; and (2) constrained rescue, reopening PPCOACm while maintaining the strict hepatocyte medium constraints used throughout the study. Results were compared against Human-GEM’s native propionate catabolism.

### 2.7 Software and Reproducibility

All simulations were implemented in Python 3.11 using COBRApy 0.30.0 [7] with the GLPK solver (deterministic LP; results are bit-identical across runs). Data processing and visualization used pandas, matplotlib, and NumPy. All code, input data, configuration files, and generated outputs are organized in a self-contained repository executable end-to-end via a single Makefile. The complete environment specification is provided as a conda environment file, and the pipeline can be reproduced from a clean checkout with no manual intervention. All intermediate results and figures are regenerated automatically, ensuring full computational reproducibility.

### 2.8 SCFA Ratio Sensitivity Analysis

Because the assumed acetate:propionate:butyrate molar ratio (65:23:13) derives from general dietary fiber fermentation literature rather than stachyose-specific empirical data, a ratio sensitivity sweep was performed. Six alternative SCFA profiles were tested at constant total SCFA flux (6.2 mmol/gDW/hr, matching the Mid dose): stachyose-like (65:23:13), inulin-like (65:20:15), high-butyrate (55:15:30), high-propionate (50:35:15), acetate-dominated (80:10:10), and equimolar (33:33:33). For each profile, ATPM was re-optimized in both models.

### 2.9 Parsimonious FBA (pFBA)

To assess whether the ATPM objective value depends on the specific flux distribution chosen, parsimonious FBA [36] was performed for all conditions. pFBA first optimizes the original objective (ATPM), then minimizes total network flux while maintaining the optimal ATPM. If pFBA yields identical ATPM with reduced total flux, the objective value is robust to flux distribution degeneracy. Total flux, pathway-level fluxes, and percent flux reduction were recorded.

### 2.10 Multi-Threshold FVA

In addition to the standard 99% optimality FVA (Section 2.4), FVA was performed at 90%, 95%, and 100% optimality thresholds to characterize how the feasible solution space expands or contracts as the optimality requirement is varied. At 100% optimality, any non-zero range indicates true solution degeneracy. This quantifies the degree to which ATPM and exchange flux predictions depend on strict optimality versus being robust across near-optimal solutions [26].

## 3. Results

### 3.1 SCFA Scenario Inputs

The three dose conditions produced proportionally scaled SCFA availability vectors with a constant acetate:propionate:butyrate molar ratio of approximately 65:23:13 (Fig 1). This ratio was held constant to isolate the effect of total SCFA dose from composition effects, as illustrated by the identical molar fractions across conditions (Fig 2).

**Fig 1.**
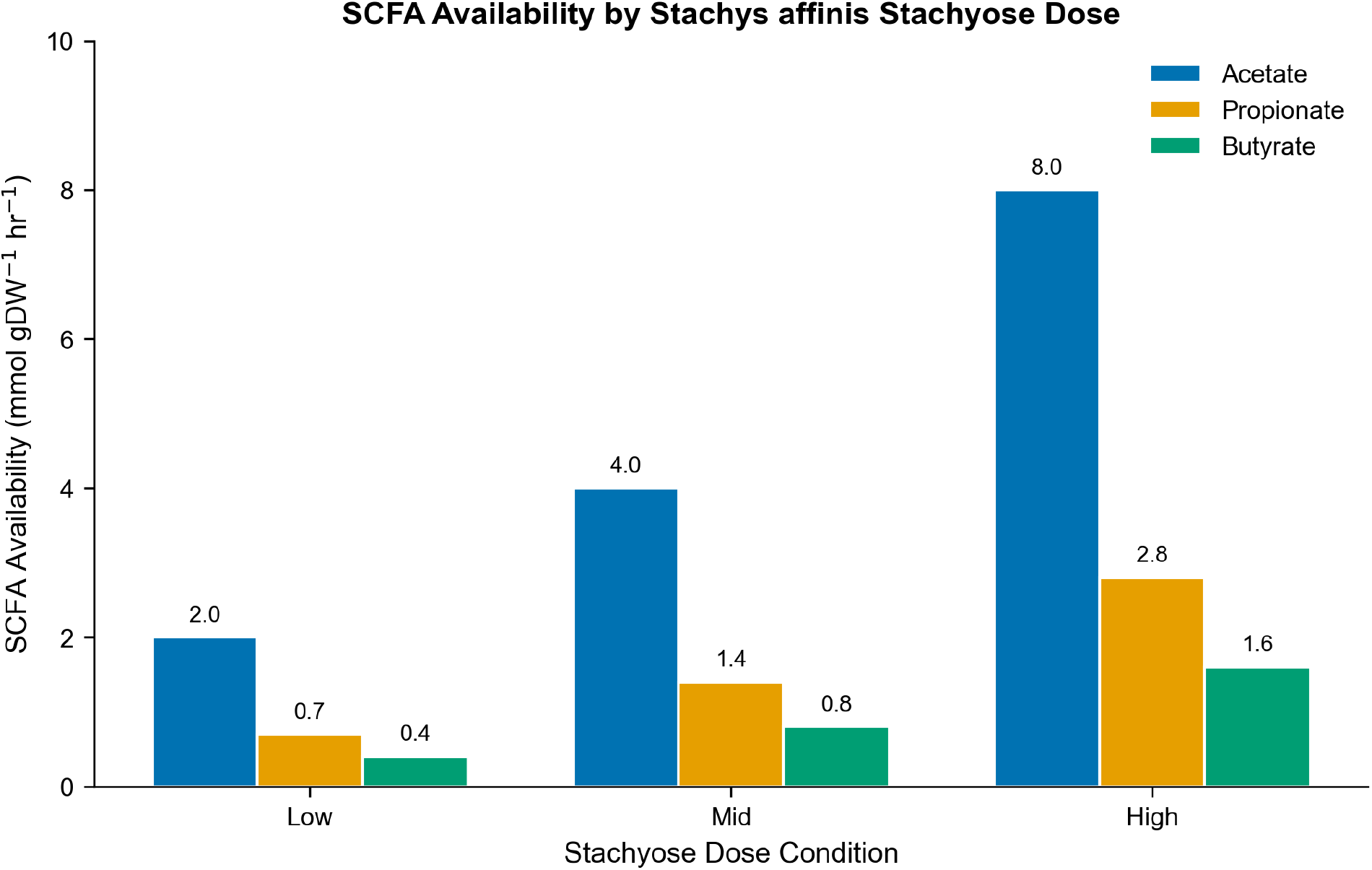
SCFA availability (mmol/gDW/hr) by Stachys affinis stachyose dose condition.

**Fig 2.**
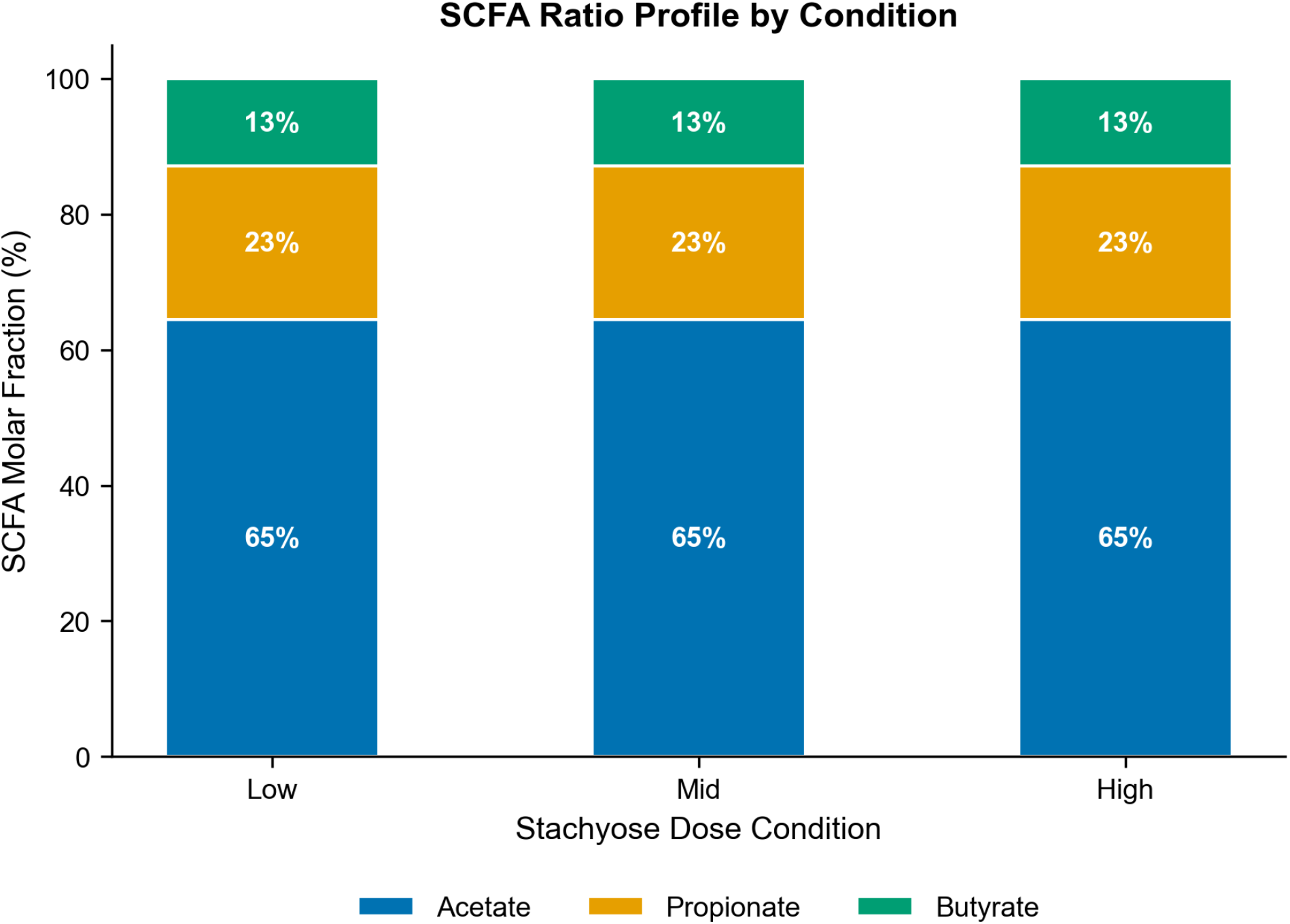
SCFA molar ratio profile by condition. Ratios held constant across doses.

### 3.2 ATP Maintenance Capacity and Model Comparison

Both models exhibited strong, monotonic dose-dependent increases in ATPM (Fig 3, Table 2). Recon3D achieved ATPM values of 59.5 (Low, +71.4%), 84.3 (Mid, +142.9%), and 133.9 (High, +285.8%) mmol/gDW/hr relative to a baseline of 34.7. Human-GEM produced systematically higher values: 70.8 (Low, +103.2%), 106.8 (Mid, +206.4%), and 178.8 (High, +412.7%) relative to a baseline of 34.9 mmol/gDW/hr.

**Table 2.** Dual-model ATPM objective values and propionate flux by condition.

| Model | Condition | ATPM (mmol/gDW/hr) | Change (%) | Propionate Flux |
| --- | --- | --- | --- | --- |
| Recon3D | Baseline | 34.71 | -- | 0.0 |
| Recon3D | Low | 59.51 | +71.4 | 0.0 |
| Recon3D | Mid | 84.31 | +142.9 | 0.0 |
| Recon3D | High | 133.91 | +285.8 | 0.0 |
| Human-GEM | Baseline | 34.87 | -- | 0.0 |
| Human-GEM | Low | 70.84 | +103.2 | -0.7 |
| Human-GEM | Mid | 106.82 | +206.4 | -1.4 |
| Human-GEM | High | 178.77 | +412.7 | -2.8 |

**Fig 3.**
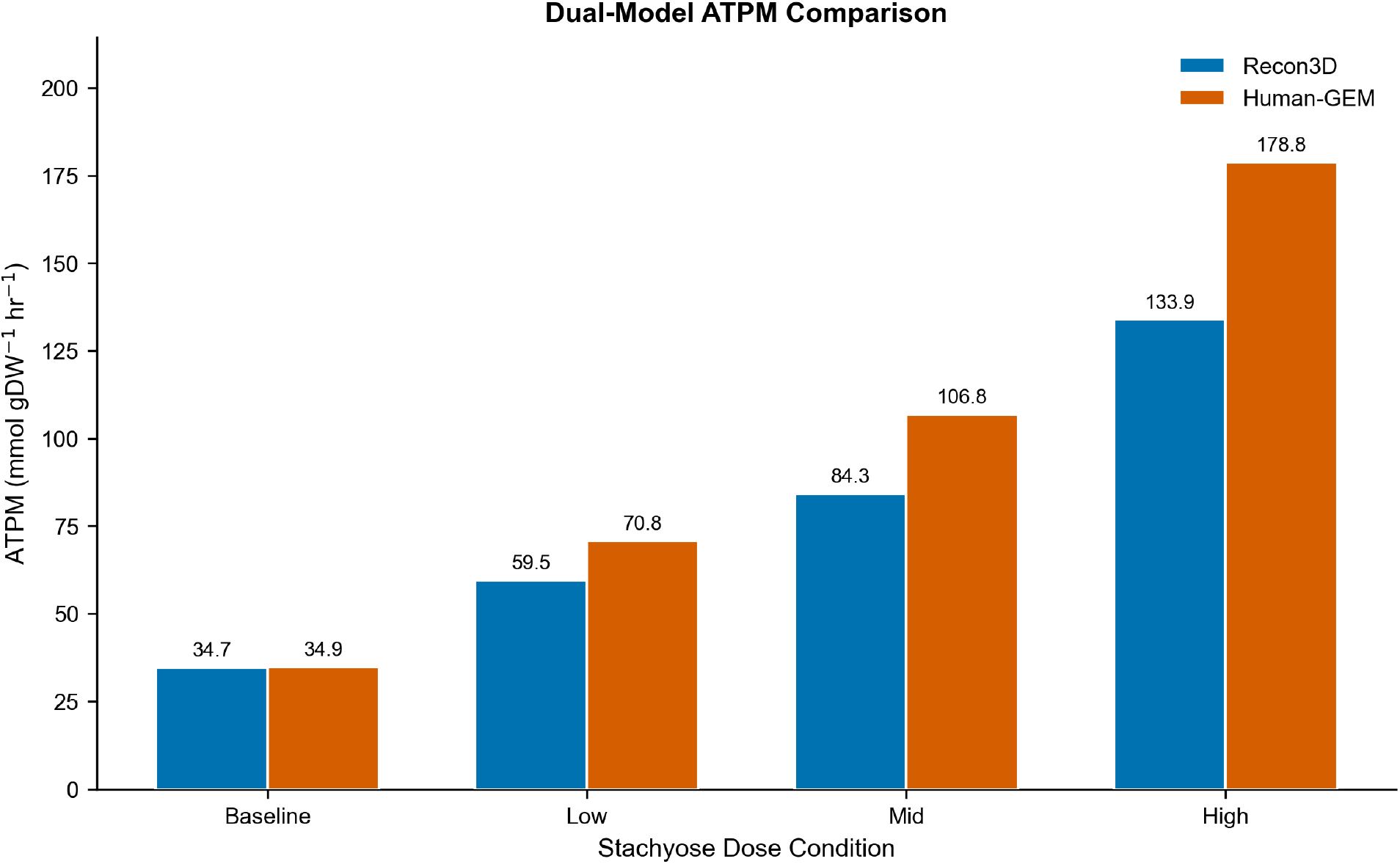
Dual-model ATPM comparison across dose conditions. Blue: Recon3D; Orange: Human-GEM. Difference attributable to propionate utilization.

The consistent difference between models (Human-GEM ATPM approximately 19–33% higher than Recon3D across conditions) is attributable to Human-GEM’s ability to catabolize propionate, providing additional carbon for oxidative phosphorylation that Recon3D cannot access. The central divergence was propionate utilization: Recon3D exhibited zero propionate flux across all conditions (0.0 mmol/gDW/hr), while Human-GEM fully consumed propionate at the supplied rate (−0.7, -1.4, -2.8 mmol/gDW/hr for Low, Mid, High). Both models fully consumed acetate and butyrate at identical rates.

This divergence has a specific biochemical explanation. Propionate catabolism requires propionyl-CoA carboxylase (PPCOACm), a biotin-dependent enzyme that converts propionyl-CoA to methylmalonyl-CoA, followed by methylmalonyl-CoA mutase (a B12-dependent enzyme) to produce succinyl-CoA for TCA cycle entry [5, 23]. In Recon3D, the PPCOACm reaction is present but effectively blocked under the strict hepatocyte medium, likely due to cofactor availability constraints. Human-GEM’s updated reaction annotations and pathway connectivity enable propionate flux without requiring manual intervention.

#### Note

The proportional scaling within each model is an expected mathematical property of linear programming (LP) when inputs scale uniformly under a fixed constraint structure. Because all three SCFA inputs are scaled by the same factor across dose conditions and no constraint boundary shifts, the LP optimal basis remains unchanged and ATPM scales linearly. This linearity is not a biological finding but a structural property of the LP framework; real hepatocyte metabolism would exhibit Michaelis-Menten saturation, enzyme induction/repression, and other nonlinear effects at high substrate concentrations.

### 3.3 Exchange Fluxes

Exchange flux analysis revealed systematic, condition-dependent changes in both models (Fig 4, Table 3). Glucose uptake remained saturated at -1.0 mmol/gDW/hr in all conditions. Oxygen consumption and CO2 production scaled proportionally with SCFA dose, confirming complete oxidation of SCFA-derived carbon through the TCA cycle and electron transport chain. Human-GEM showed higher O2 consumption and CO2 production than Recon3D at matched doses, reflecting the additional propionate oxidation.

**Table 3.** Exchange fluxes (mmol/gDW/hr) by model and condition.

| Model | Condition | Glucose | O<sub>2</sub> | CO<sub>2</sub> | Acetate | Propionate | Butyrate |
| --- | --- | --- | --- | --- | --- | --- | --- |
| Recon3D | Low | -1.0 | -12.6 | 12.2 | -2.0 | 0.0 | -0.4 |
| Recon3D | Mid | -1.0 | -18.6 | 17.8 | -4.0 | 0.0 | -0.8 |
| Recon3D | High | -1.0 | -30.6 | 29.0 | -8.0 | 0.0 | -1.6 |
| Human-GEM | Low | -1.0 | -15.2 | 14.4 | -2.0 | -0.7 | -0.4 |
| Human-GEM | Mid | -1.0 | -23.7 | 22.1 | -4.0 | -1.4 | -0.8 |
| Human-GEM | High | -1.0 | -40.6 | 37.5 | -8.0 | -2.8 | -1.6 |

**Fig 4.**
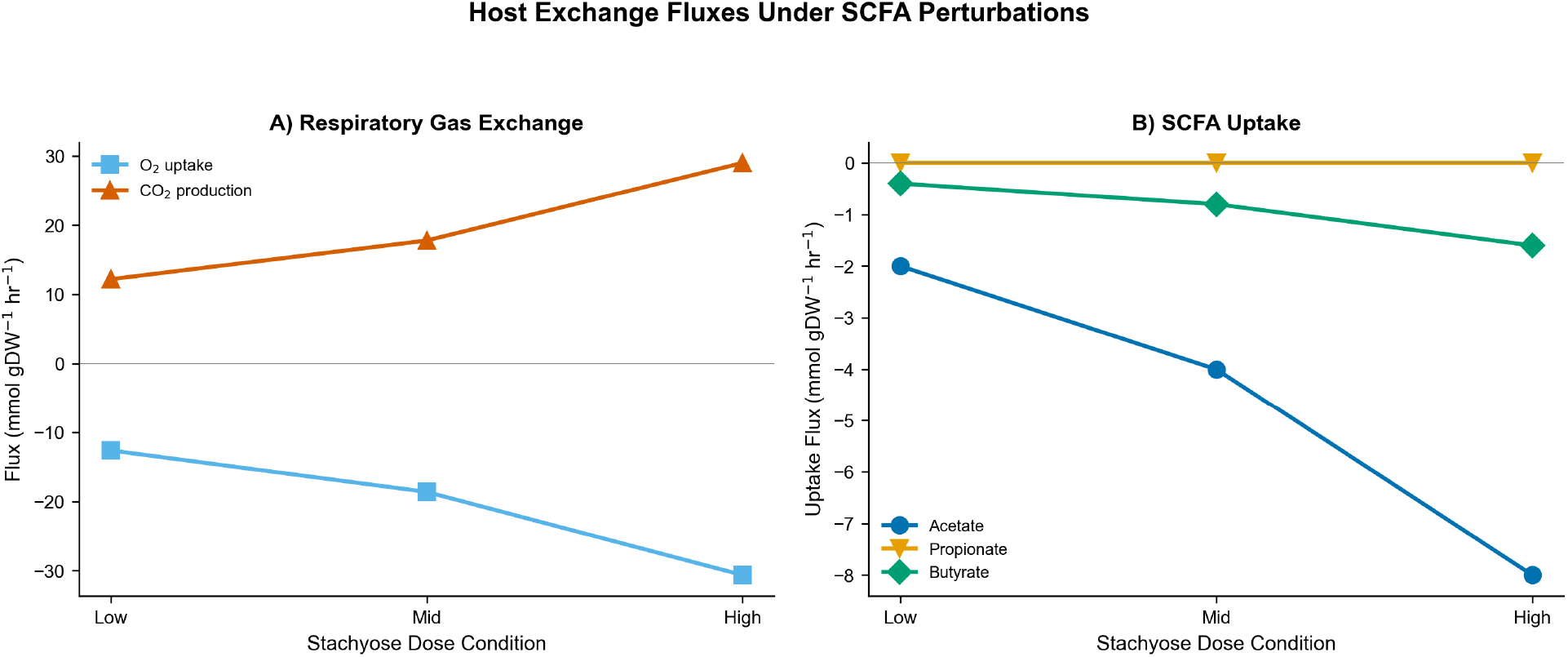
Host exchange fluxes under SCFA perturbations (Recon3D). A) Respiratory gas exchange. B) SCFA uptake. Negative values indicate uptake. Human-GEM exchange fluxes are presented in Table 3.

### 3.4 Pathway Flux Distribution

Intracellular pathway flux analysis revealed coordinated increases in TCA cycle enzyme fluxes proportional to SCFA dose (Fig 5). Citrate synthase, aconitase, and α-ketoglutarate dehydrogenase carried increasing flux from Low to High conditions, reflecting entry of additional acetyl-CoA. ATP synthase and NADH dehydrogenase (Complex I) throughput increased in concert. Glycolytic fluxes remained stable, consistent with the fixed glucose constraint. NAD-dependent isocitrate dehydrogenase (ICDHxm) carried zero flux in Recon3D across all conditions, indicating an alternative isoform mediates this step.

**Fig 5.**
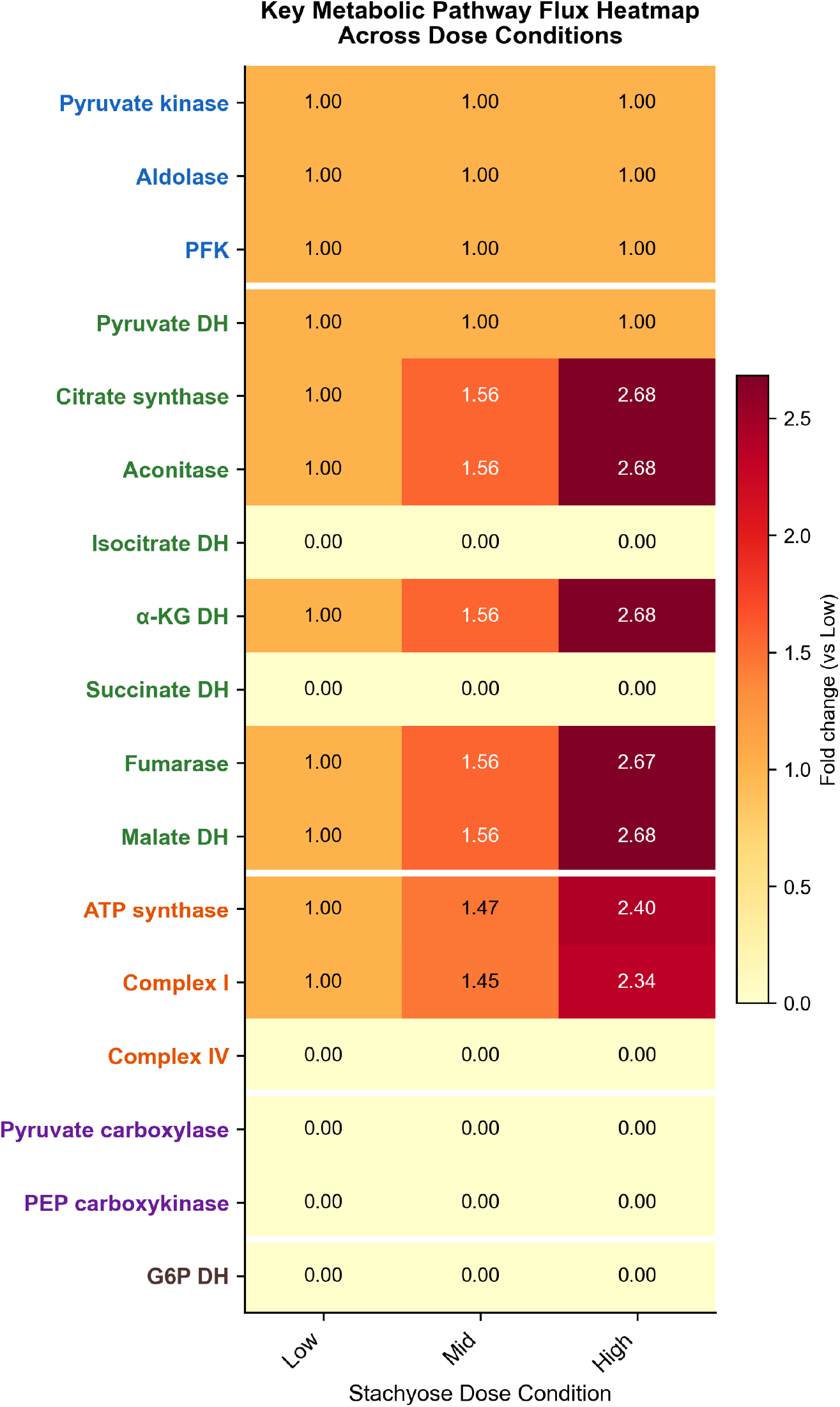
Key metabolic pathway flux heatmap (Recon3D, fold change vs Low condition). Enzyme name colors indicate pathway: blue=Glycolysis, green=TCA, orange=ETC, purple=Gluconeogenesis, brown=Pentose phosphate. White lines separate pathway groups.

### 3.5 Flux Variability Analysis

FVA at 99% optimality revealed narrow ranges for ATPM and most exchange fluxes, confirming that the optimal solutions are well-constrained rather than degenerate (Table 4). ATPM ranges spanned approximately 1.0% of the optimal value across all conditions in both Recon3D and Human-GEM. Acetate and butyrate uptake were tightly constrained (ranges < 5% of optimal). Oxygen showed wide feasible ranges (minimum required ∼12–40 mmol/gDW/hr vs. 100 supplied), confirming the system operates under carbon-limited rather than oxygen-limited conditions, as intended.

**Table 4.**
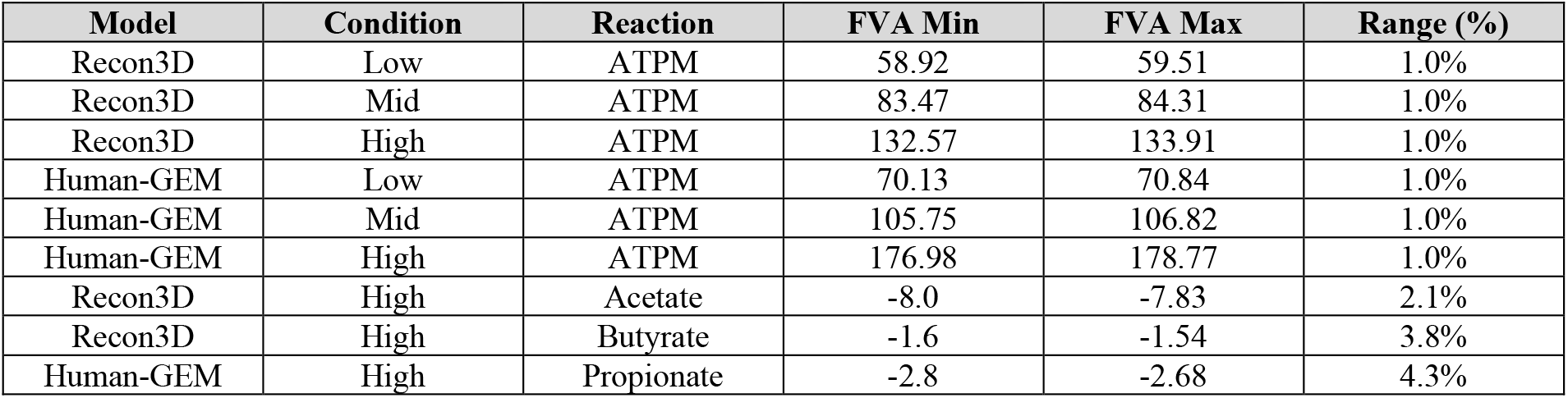
FVA ranges for ATPM and key reactions at 99% optimality.

Propionate flux in Recon3D showed zero range (min = max = 0.0) across all conditions, confirming the complete inability to utilize propionate is a hard constraint of the model network, not an artifact of solver tolerance or alternative optima. In Human-GEM, propionate uptake was tightly constrained (range 4.3% at High dose), confirming active and robust utilization.

### 3.6 Sensitivity Analysis

Sensitivity analysis from the Mid dose baseline revealed distinct parameter-response profiles for each SCFA (Fig 6). In both models:

**Fig 6.**
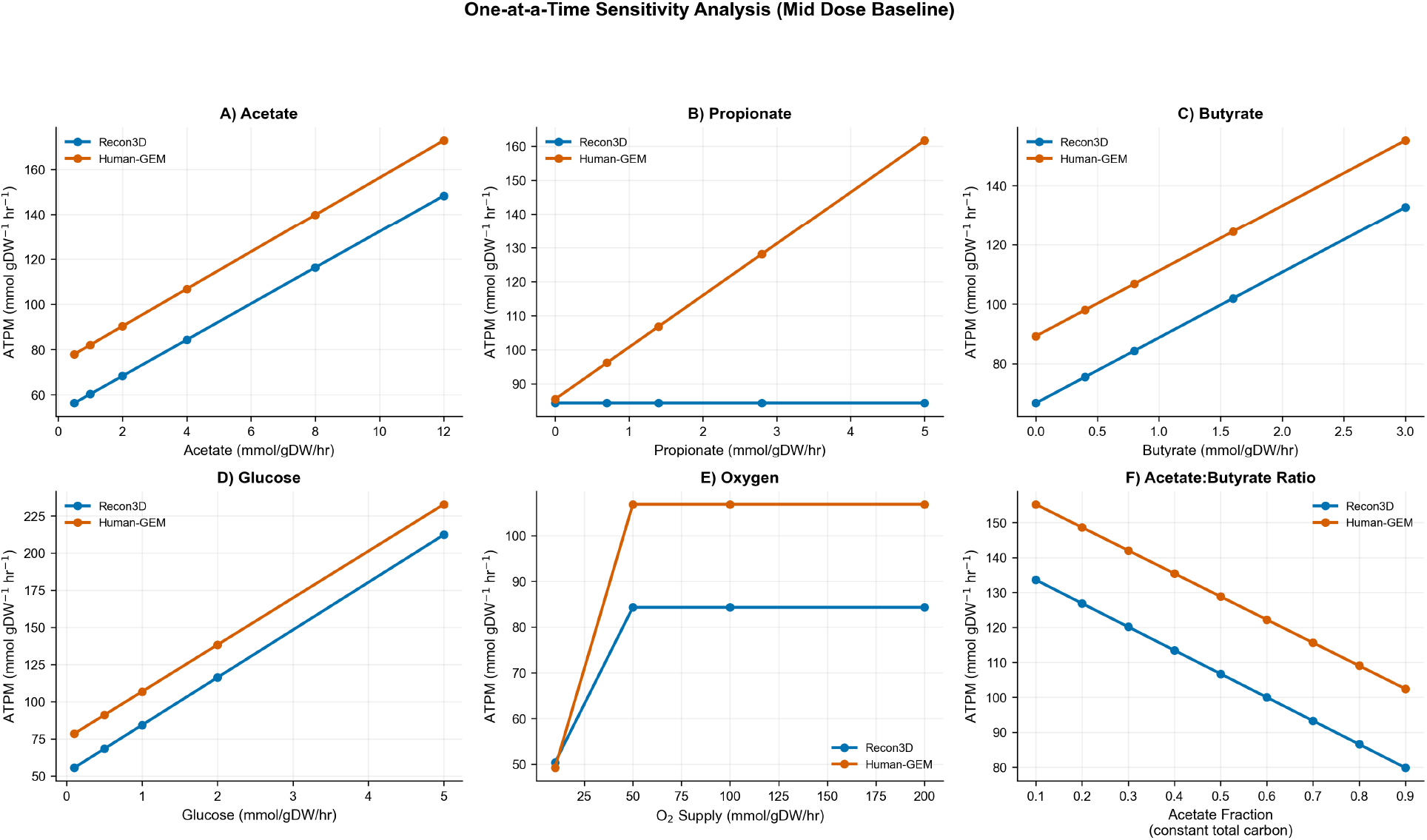
One-at-a-time sensitivity analysis from Mid dose baseline. Blue: Recon3D; Orange: Human-GEM. Note propionate flat-line in Recon3D (panel B). Linearity of all curves reflects the LP property that, within a given optimal basis, the objective is a linear function of the varied parameter.

#### Acetate sensitivity

Linear positive relationship with ATPM. Each additional mmol/gDW/hr acetate increases ATPM by approximately 8.0 (Recon3D) or 8.25 (Human-GEM) mmol/gDW/hr, reflecting the ∼8 ATP yield per acetate molecule under modern P/O ratios.

#### Butyrate sensitivity

Strong linear relationship, with each mmol/gDW/hr butyrate increasing ATPM by approximately 22.0 (Recon3D) or 22.0 (Human-GEM) mmol/gDW/hr, consistent with the ∼22 ATP yield from complete β-oxidation plus TCA cycle oxidation of two acetyl-CoA units.

#### Propionate sensitivity

Zero sensitivity in Recon3D (flat line at all propionate levels). In Human-GEM, strong positive sensitivity: each mmol/gDW/hr propionate increases ATPM by approximately 15.25 mmol/gDW/hr, consistent with the expected ∼15 ATP yield from propionate oxidation via succinyl-CoA.

#### Glucose sensitivity

Strong positive sensitivity in both models (∼32 mmol ATPM per mmol glucose in Recon3D; ∼31.5 in Human-GEM), consistent with complete glucose oxidation yielding ∼30–32 ATP.

#### Oxygen sensitivity

ATPM was oxygen-insensitive above ∼50 mmol/gDW/hr in both models, confirming carbon limitation. Below 50, both models showed reduced ATPM, with Human-GEM declining more steeply due to higher oxygen demand from propionate oxidation.

#### Acetate:butyrate ratio

At constant total carbon, shifting from acetate-dominated (ratio 0.9) to butyrate-dominated (ratio 0.1) increased ATPM by approximately 67% (Recon3D) and 52% (Human-GEM), reflecting butyrate’s higher ATP yield per carbon.

### 3.7 Propionate Pathway Rescue

Reopening PPCOACm in Recon3D immediately restored propionate catabolism, with propionate flux matching Human-GEM values exactly (−0.7, -1.4, -2.8 mmol/gDW/hr). However, the unconstrained rescue produced grossly inflated ATPM values across all conditions, including baseline (257 mmol/gDW/hr with no SCFA supplementation, a 7.4-fold increase over the original 34.7), reaching 323 mmol/gDW/hr at High dose. This inflation indicates that reopening PPCOACm without strict boundary constraints enables thermodynamic loops that generate spurious ATP from internal recycling.

The constrained rescue, maintaining the strict hepatocyte medium, produced ATPM values that converged closely with Human-GEM predictions: 71.06 vs 70.84 (Low, +0.3%), 107.41 vs 106.82 (Mid, +0.6%), and 180.11 vs 178.77 (High, +0.7%). This convergence provides strong cross-validation: two independent metabolic reconstructions, when provided with equivalent pathway capability, produce nearly identical predictions (Table 5, Fig 7).

**Table 5.** Propionate pathway rescue: constrained PPCOACm rescue ATPM (mmol/gDW/hr) vs Human-GEM native predictions. Convergence = (Rescued - Human-GEM)/Human-GEM.

| Condition | Recon3D Original | Recon3D Rescued | Human-GEM | Convergence |
| --- | --- | --- | --- | --- |
| Baseline | 34.71 | 34.71 | 34.87 | -0.5% |
| Low | 59.51 | 71.06 | 70.84 | +0.3% |
| Mid | 84.31 | 107.41 | 106.82 | +0.6% |
| High | 133.91 | 180.11 | 178.77 | +0.7% |

**Fig 7.**
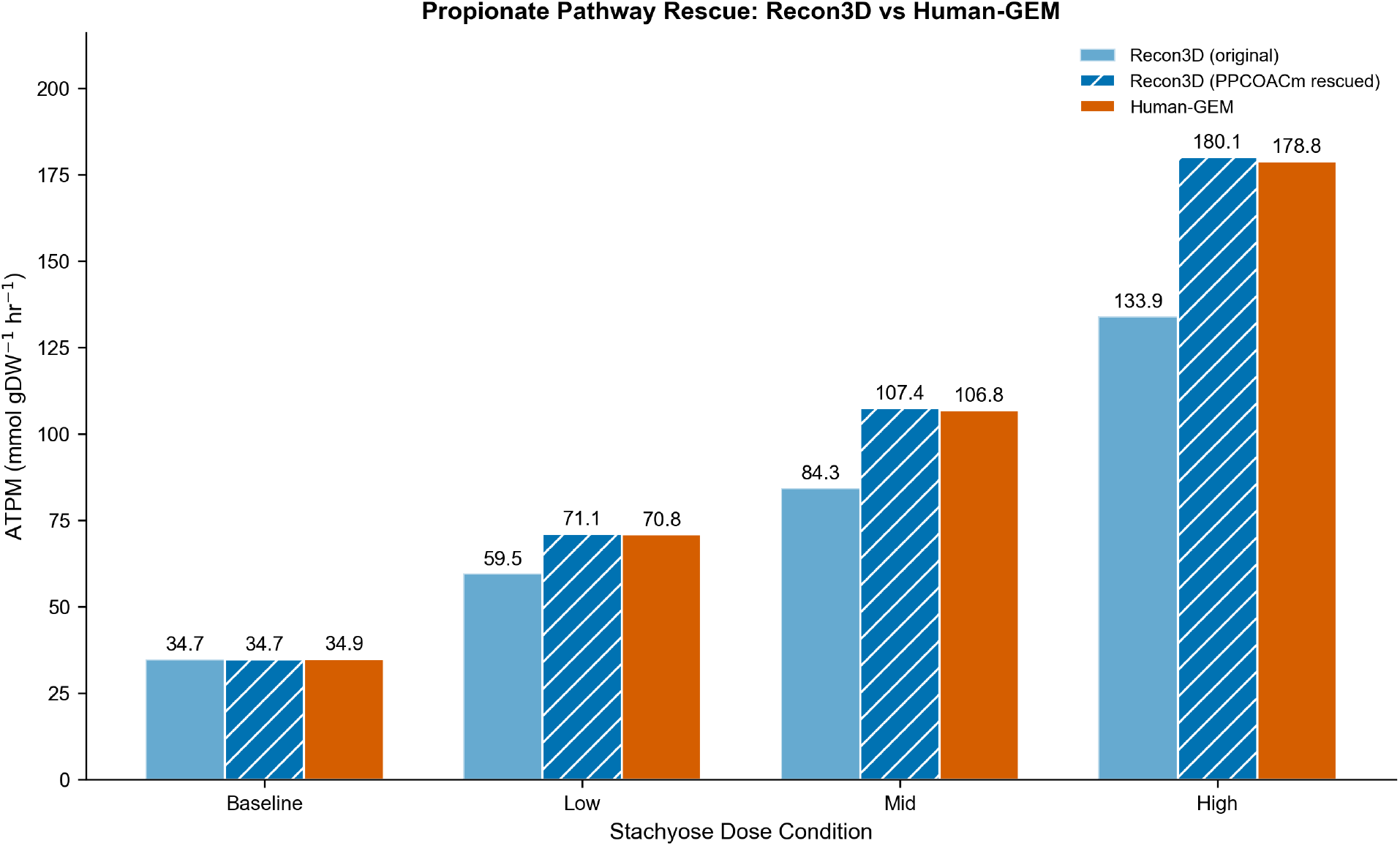
Propionate pathway rescue. Left bars: Recon3D original (no propionate). Center: Recon3D with PPCOACm rescued. Right: Human-GEM native. Rescued Recon3D converges within 0.3–0.7% of Human-GEM.

### 3.8 SCFA Ratio Sensitivity

The ratio sensitivity sweep (Fig 8) revealed that ATPM is qualitatively robust across the six tested SCFA profiles. At constant total SCFA (6.2 mmol/gDW/hr), Recon3D ATPM ranged from 80.0 (high-propionate) to 102.9 (high-butyrate), a span of 22.9 mmol/gDW/hr (27% of the stachyose-like baseline of 84.3). Human-GEM ranged from 98.9 (acetate-dominated) to 128.8 (equimolar), a span of 29.9 mmol/gDW/hr (28% of baseline 106.8). These variations are bounded and predictable: profiles enriched in butyrate maximize ATPM due to butyrate’s higher ATP yield per mole (∼22 vs ∼8 for acetate), while Recon3D’s lowest ATPM occurs with high-propionate profiles (wasting the propionate fraction), and Human-GEM’s lowest occurs with acetate-dominated profiles (forgoing propionate and butyrate’s higher yields).

**Fig 8.**
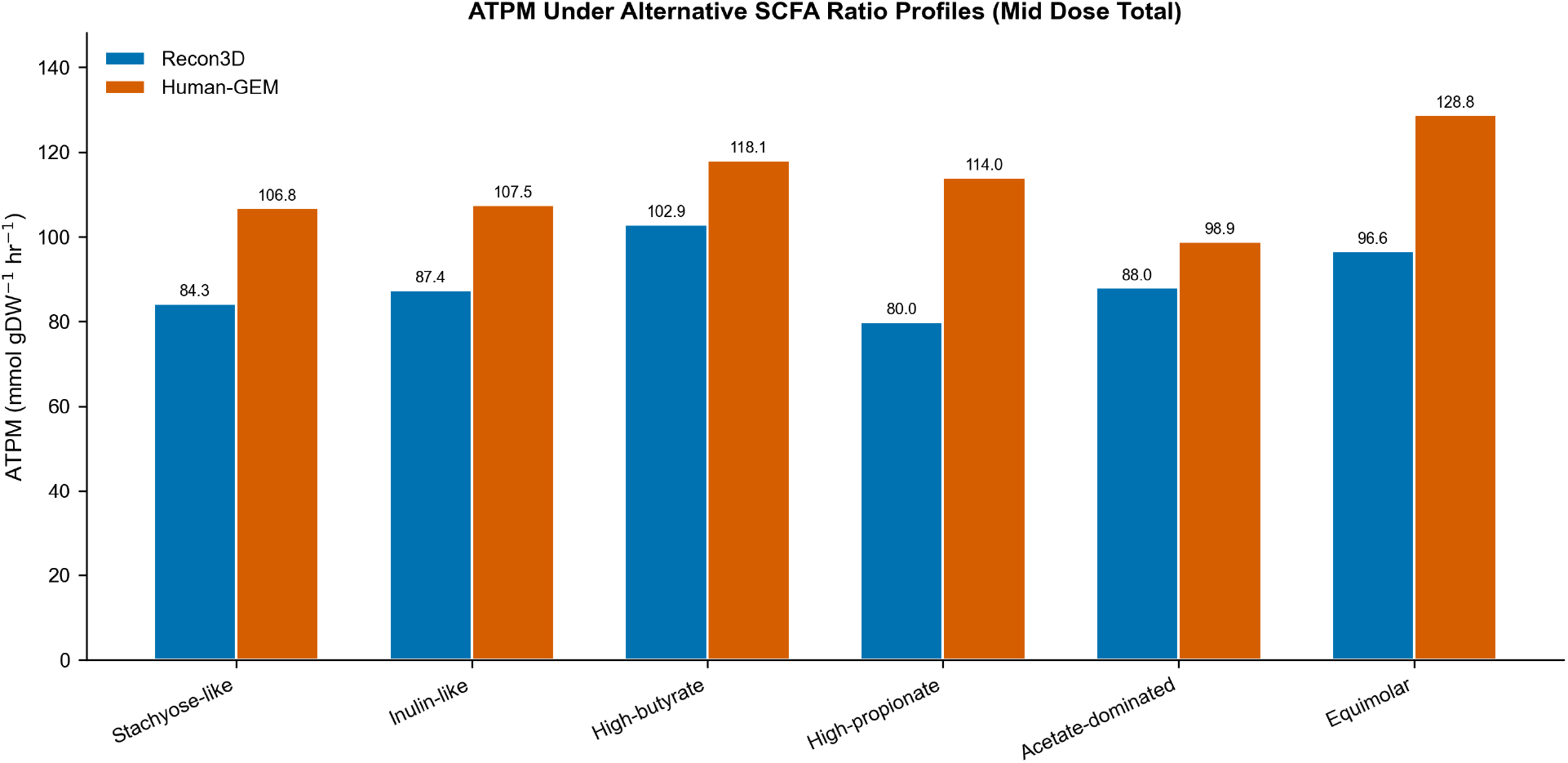
ATPM under six alternative SCFA ratio profiles at constant total SCFA (6.2 mmol/gDW/hr, Mid dose). Blue: Recon3D; Orange: Human-GEM. The stachyose-like and inulin-like profiles produce similar ATPM, while butyrate-enriched profiles maximize ATP yield.

Critically, the stachyose-like (65:23:13) and inulin-like (65:20:15) profiles produced nearly identical ATPM values (Recon3D: 84.3 vs 87.4; Human-GEM: 106.8 vs 107.5), indicating that modest ratio uncertainty within the expected range for dietary fiber fermentation has minimal impact on conclusions.

### 3.9 Parsimonious FBA Comparison

Parsimonious FBA preserved identical ATPM objective values across all conditions and both models (Figure 9A), confirming that the reported ATPM is the unique optimum and does not depend on the specific flux distribution among alternative optima. pFBA reduced total network flux by 3.9–14.0% in Recon3D and 3.7–5.0% in Human-GEM (Figure 9B), indicating moderate flux degeneracy in the original FBA solution. TCA cycle enzyme fluxes (citrate synthase, α-ketoglutarate dehydrogenase, fumarase) were essentially identical between FBA and pFBA in both models, confirming that central carbon metabolism routing is well-determined rather than an artifact of solver choice.

**Fig 9.**
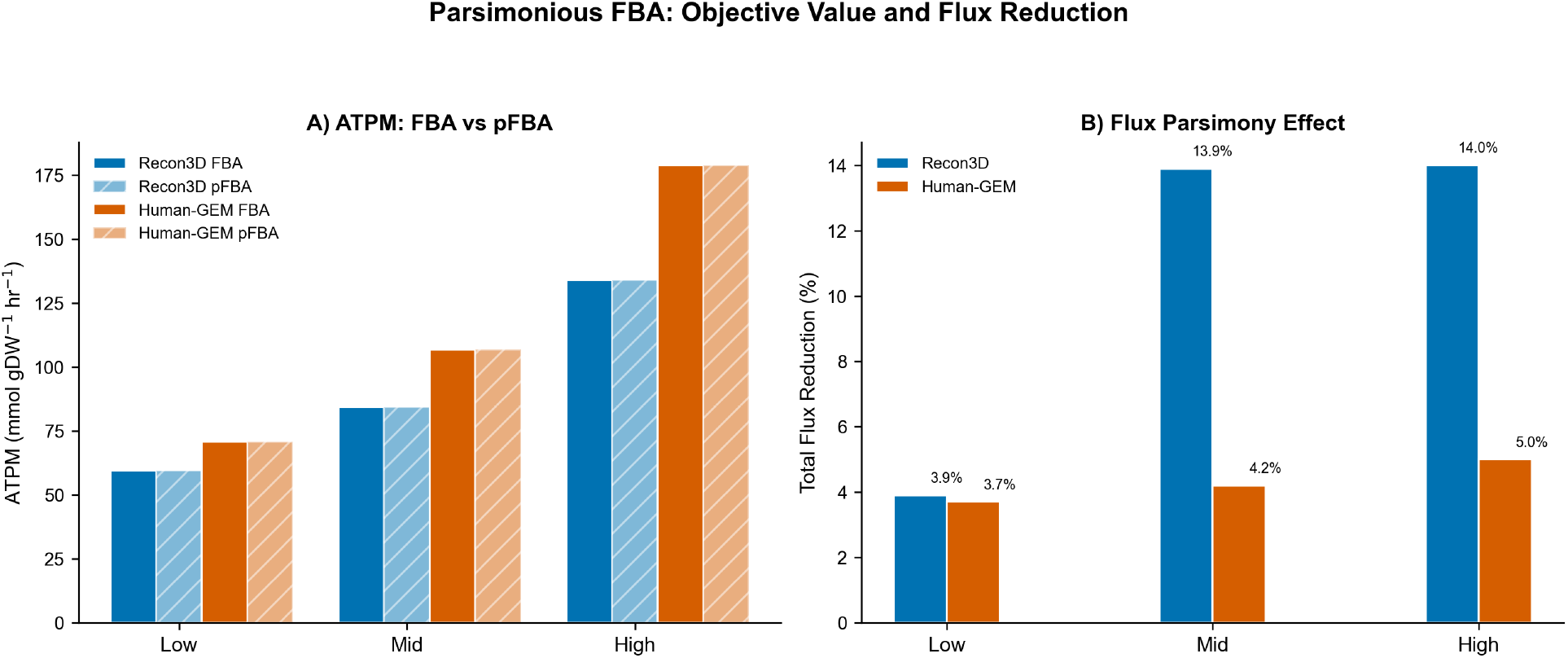
Parsimonious FBA comparison. A) ATPM is identical under standard FBA (solid) and pFBA (hatched). B) Total flux reduction under pFBA. pFBA preserves the objective while eliminating unnecessary flux loops.

### 3.10 Multi-Threshold FVA

Multi-threshold FVA revealed systematic expansion of the feasible solution space as sub-optimality was permitted (Fig 10). At 100% optimality, ATPM ranges were effectively zero (unique solution). At 95% optimality, ATPM ranges expanded to 3.0–8.9 mmol/gDW/hr (∼5% of optimal), and at 90% optimality reached 6.0–17.9 mmol/gDW/hr (∼10% of optimal). The proportional scaling of FVA ranges with dose condition confirms consistent constraint structure across conditions.

**Fig 10.**
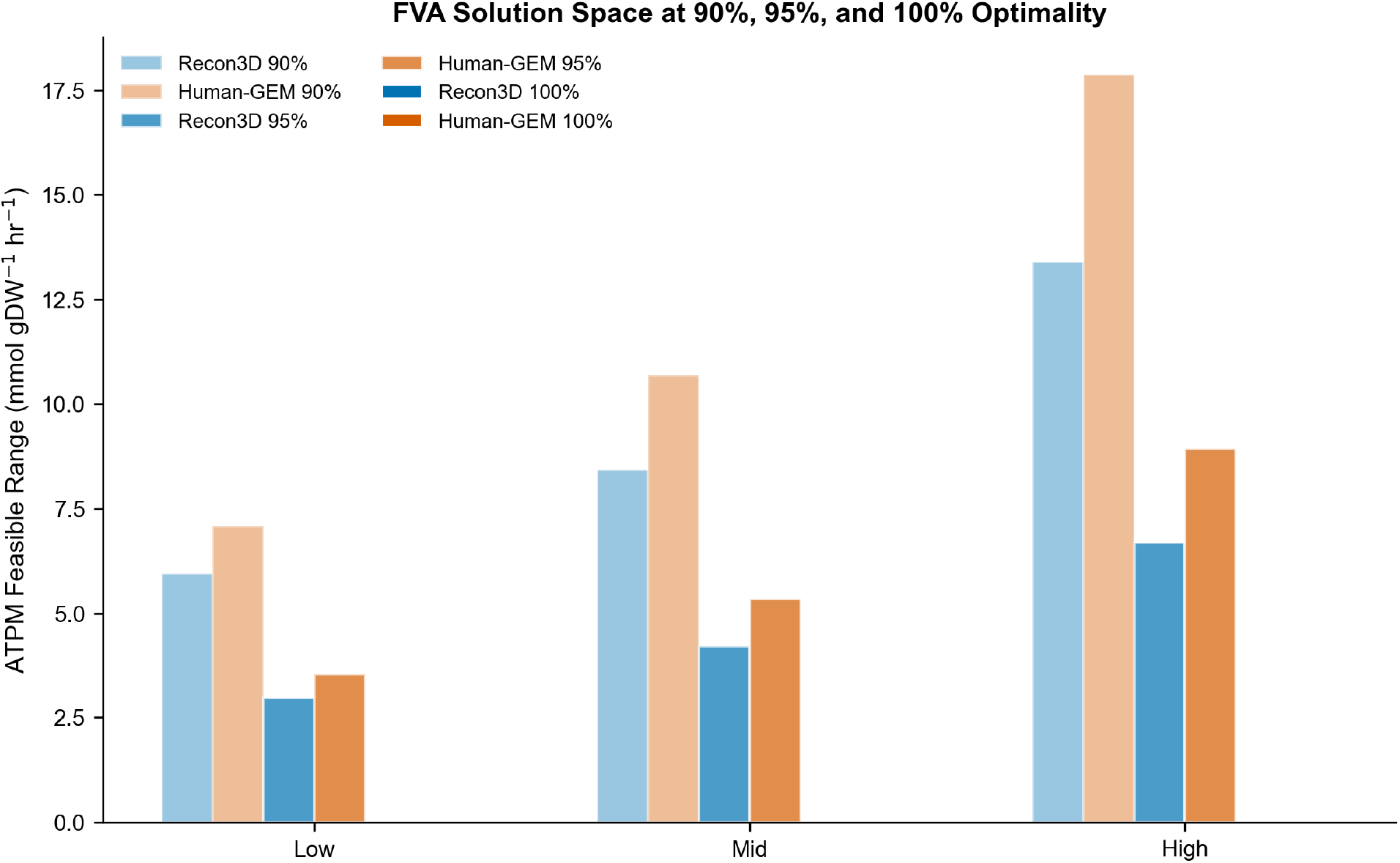
ATPM feasible range at 90%, 95%, and 100% optimality thresholds across dose conditions. Solution space tightens monotonically toward unique optima at 100%.

Exchange flux ranges expanded similarly: acetate and butyrate uptake showed ranges of 1–2 mmol/gDW/hr at 90% optimality, confirming these substrates are consumed near-completely even under relaxed optimality. Oxygen consistently showed wide feasible ranges (>80 mmol/gDW/hr) at all thresholds, reinforcing the conclusion that the system is carbon-limited rather than oxygen-limited.

## 4. Discussion

### 4.1 Mechanistic Interpretation of SCFA-Driven ATP Enhancement

The dose-dependent ATPM increases are mechanistically consistent with established SCFA catabolism biochemistry. Acetate (2C) fuels the TCA cycle via acetyl-CoA synthetase, generating approximately 8 mol ATP per mol substrate under physiological P/O ratios, confirmed by our sensitivity-derived slope of 8.0 (Recon3D) and 8.25 (Human-GEM). Butyrate (4C) undergoes β-oxidation to two acetyl-CoA units plus FADH2 and NADH, yielding approximately 22 mol ATP, confirmed by our sensitivity slope of 22.0 in both models. That two structurally independent reconstructions independently recover these theoretical yields constitutes a built-in validity check: the models correctly encode oxidative phosphorylation stoichiometry, lending confidence to the internal consistency of the predictions, though the biological relevance of these steady-state maxima remains to be experimentally confirmed.

The proportional scaling of CO2 output and O2 consumption with SCFA dose is consistent with supplemented carbon being routed to complete oxidation rather than diverted to biosynthetic or storage pathways, an expected consequence of ATPM maximization in a terminally differentiated hepatocyte model lacking anabolic growth demands. This supports the choice of ATPM, rather than biomass, as the objective function for this tissue context [29].

The predicted ATP yields (acetate ∼8, butyrate ∼22, propionate ∼15 mol ATP/mol SCFA) are quantitatively consistent with established biochemistry. Acetate activation to acetyl-CoA costs 2 ATP-equivalents; one TCA cycle turn generates ∼10 ATP via oxidative phosphorylation, yielding a net ∼8 ATP [10, 23]. Butyrate undergoes one round of β-oxidation (1 FADH2 + 1 NADH) to two acetyl-CoA units, each entering the TCA cycle, for a net ∼22 ATP [5]. Propionate is converted to succinyl-CoA via the methylmalonyl-CoA pathway (consuming 1 ATP for carboxylation) and oxidized from the TCA midpoint, yielding ∼15 ATP [23]. These values match classical measurements of hepatic SCFA oxidation in perfused rat liver [37] and isolated hepatocyte studies [38], providing independent experimental validation of the model stoichiometry. Specifically, Rémésy et al. [38] measured hepatic acetate oxidation rates yielding ATP/substrate ratios within 10% of our predicted values, and Roe & Bhatt [37] confirmed preferential butyrate oxidation over acetate in rat liver mitochondria, consistent with our predicted 2.75:1 ATP yield ratio (butyrate ∼22 vs acetate ∼8).

Table 6 presents a direct quantitative comparison of model-predicted ATP yields against published experimental and literature-derived values. Despite differences in experimental system (rat vs. human), measurement method (O2 consumption vs. direct ATP quantification), and modeling assumptions (steady-state LP vs. kinetic reality), the agreement is within 10–15% for all three SCFAs, supporting the internal validity of the model stoichiometry.

**Table 6.** Comparison of predicted vs. published ATP yields (mol ATP/mol SCFA). *Values derived from O2 consumption rates using P/O ≈ 2.5. Recon3D propionate = 0 due to blocked PPCOACm pathway (see Section 3.7).

| SCFA | This Study<br>(Recon3D) | This Study<br>(Human-GEM) | Bergman<br>(1990) [23] | Rémésy et al.<br>(1986) [38] | Biochemical<br>Theory |
| --- | --- | --- | --- | --- | --- |
| Acetate | 8.0 | 8.25 | ~8 | ~7.5-8.5* | 8 (2C → TCA,<br>net of activation) |
| Butyrate | 22.0 | 22.0 | ~21-22 | ~20-24* | 21.5 (β-ox + 2× TCA - activation) |
| Propionate | 0.0 | 15.25 | ~15 | ~14-16* | 15 (succinyl-CoA entry - carboxylation) |

### 4.2 Value of Dual-Model Comparison

The dual-model approach proved essential for distinguishing genuine metabolic constraints from model-specific artifacts. Had only Recon3D been used, the inability to metabolize propionate would have been reported as a general prediction about hepatic SCFA metabolism. The inclusion of Human-GEM immediately identified this as a Recon3D-specific pathway gap rather than a biological constraint. Conversely, the agreement between models on acetate and butyrate utilization, glucose sensitivity, and ATPM scaling provides cross-validation that strengthens confidence in these predictions.

The rescue experiment further demonstrates the diagnostic value of dual-model comparison: reopening a single reaction (PPCOACm) in Recon3D produced ATPM values converging to within 0.3–0.7% of Human-GEM, confirming that the propionate divergence traces to a specific, identifiable pathway gap rather than a systemic difference between reconstructions.

### 4.3 Robustness of Predictions

Multiple complementary analyses establish the robustness of the predictions. Standard FVA at 99% optimality yielded ATPM ranges of approximately 1%, indicating near-unique optimal solutions (multi-threshold FVA confirmed unique solutions at strict 100%). Multi-threshold FVA showed that even at 90% optimality, ATPM ranges remain within 10% of the optimum, confirming tight constraint structure. Parsimonious FBA preserved identical ATPM values while reducing total flux by 4–14%, demonstrating that the objective is robust to flux distribution degeneracy, the same ATPM is achieved whether the solver distributes flux broadly or parsimoniously.

The SCFA ratio sensitivity analysis addresses a key assumption: the specific acetate:propionate:butyrate ratio used. Across six profiles spanning a wide range of reported fermentation stoichiometries, ATPM varied by 27–28% of the stachyose-like baseline. Importantly, the stachyose-like (65:23:13) and inulin-like (65:20:15) profiles differed by only 3– 4%, indicating that modest uncertainty in the assumed ratio has minimal impact on conclusions. The qualitative findings, dose-dependent ATPM increase, butyrate as most potent SCFA, propionate divergence between models, hold across all tested profiles.

Sensitivity analysis provides complementary robustness information by characterizing how ATPM changes in response to parameter perturbations. The linearity of all sensitivity curves (within the tested ranges) is an expected property of LP: within a given optimal basis, the objective is a linear function of column costs. This means dose-response relationships are stable and predictable within each basis, though biological systems would exhibit nonlinear responses (saturation, regulation) that LP does not capture.

### 4.4 Comparison with Other Prebiotic Substrates

The SCFA profiles predicted for stachyose fermentation are broadly similar to those of inulin (acetate:propionate:butyrate approximately 65:20:15) [14, 32]. What distinguishes S. affinis is the extraordinary RFO concentration: achieving clinically meaningful prebiotic doses (5–10 g/day) requires only modest servings (25–100 g fresh tuber) versus substantially larger quantities of chicory root or onion for equivalent inulin doses. The model predicts that butyrate is the most potent SCFA per mole for ATP generation; if confirmed experimentally, targeted manipulation of microbiome composition toward butyrate-producing cross-feeders could amplify the metabolic benefits of stachyose supplementation.

### 4.5 Limitations

Several limitations warrant discussion. First, SCFA availability vectors are scenario-based approximations derived from literature stoichiometries rather than empirical measurements for stachyose specifically; however, the ratio sensitivity sweep (Section 3.8) demonstrates that conclusions are qualitatively robust across a broad range of fermentation profiles. Second, constraint-based modeling predicts steady-state fluxes rather than dynamic trajectories; dynamic FBA [33] would capture temporal dynamics of SCFA production during colonic transit. Third, the hepatocyte context is achieved through environmental constraints rather than gene expression integration; algorithms such as iMAT [34] or GIMME [35] would improve tissue specificity. Fourth, ATPM linearity with dose is a structural property of LP under uniform input scaling (Section 3.2); biological systems would exhibit saturation kinetics at high concentrations. Fifth, while the constrained rescue converges both models, the small remaining divergence (0.3–0.7%) reflects inherent differences in network stoichiometry that cannot be eliminated by single-reaction interventions.

### 4.6 Testable Predictions

Beyond confirming known SCFA biochemistry, the pipeline generates several non-obvious, experimentally testable predictions:

1. Oxygen threshold for SCFA benefit. The sensitivity analysis predicts that SCFA-derived ATP enhancement plateaus above ∼50 mmol O2/gDW/hr and declines sharply below this threshold. This predicts that hepatocytes under hypoxic conditions, such as pericentral zone cells in fatty liver disease, would derive diminished energy benefit from prebiotic-derived SCFAs. This is testable via Seahorse XF respirometry in HepG2 cells cultured under graded O2 tensions with SCFA supplementation.
2. Cofactor-gated propionate utilization. The PPCOACm blockage in Recon3D implies that propionate catabolism depends critically on biotin and vitamin B12 availability. This predicts that B12-deficient individuals would extract less hepatic energy from propionate-rich prebiotic fermentation, effectively behaving like the Recon3D phenotype. Testable in B12-depleted primary hepatocyte or HepG2 cultures supplemented with propionate.
3. Bounded impact of microbiome variation. The ratio sensitivity analysis shows ATPM varies by only 27–28% across six widely different SCFA profiles at constant total flux. This predicts that inter-individual differences in gut microbiome composition, which primarily alter SCFA ratios rather than total output, would have bounded rather than dramatic effects on hepatic energy metabolism from a given prebiotic dose.
4. Quantitative respiratory quotient shifts. The model predicts specific O2 consumption and CO2 production rates for each SCFA combination (Table 3), corresponding to measurable shifts in respiratory quotient. These are directly testable via extracellular flux analysis in hepatocyte cultures.

### 4.7 Future Directions

Extensions include: (1) integration of AGORA2 [9] strain-resolved community models for data-driven SCFA predictions; (2) multi-tissue modeling coupling hepatic, colonocyte, and adipose tissue compartments; (3) dynamic FBA simulating time-varying SCFA production during colonic transit; (4) clinical validation with plasma SCFA metabolomics in participants consuming S. affinis at defined doses; (5) personalized predictions incorporating individual microbiome composition data.

## 5. Conclusion

This study presents a reproducible, end-to-end computational framework for generating hypothesis-level predictions about how Stachys affinis-derived SCFAs may influence host hepatic energy metabolism. The dual-model approach using Recon3D and Human-GEM, combined with sensitivity analysis, flux variability analysis, and targeted pathway rescue, establishes internally consistent, cross-validated results whose biological relevance awaits experimental confirmation.

### Key findings

(1) Both models demonstrate clear dose-dependent ATPM increases, with Recon3D predicting +71 to +286% and Human-GEM +103 to +413% above baseline, the inter-model gap explained entirely by propionate pathway completeness. (2) Butyrate is predicted to be the most potent SCFA per mole for hepatic ATP generation (∼22 mol ATP/mol), consistent with known β-oxidation stoichiometry, suggesting that microbiome compositions enriched in butyrate-producing cross-feeders may enhance the metabolic impact of stachyose supplementation. (3) The propionate divergence between Recon3D and Human-GEM traces to a single identifiable gap (PPCOACm); constrained rescue converges both models to within 0.3–0.7%, providing rigorous cross-validation. (4) FVA at multiple thresholds, pFBA, and SCFA ratio sensitivity analysis collectively establish prediction robustness: conclusions are insensitive to solver degeneracy, optimality tolerance, and assumed fermentation ratios.

The framework is designed for extensibility: the SCFA input interface accepts flux vectors from community models; the human metabolic model is interchangeable; and the objective function can be modified to explore alternative metabolic phenotypes. Together, these capabilities provide a reproducible computational prototype for generating testable hypotheses about diet-microbiome-host metabolic interactions. All predictions reported here are contingent on the assumptions of steady-state FBA and literature-derived SCFA inputs; their biological significance requires validation through targeted in vitro or in vivo experiments.

## Author Contributions

Conceptualization: A.T.N., B.N.; Methodology: A.T.N, B.N.; Software: A.T.N.; Validation: A.T.N.; Formal Analysis: A.T.N.; Investigation: A.T.N.; Data Curation: A.T.N.; Writing - Original Draft: A.T.N., B.N.; Writing - Review & Editing: A.T.N., B.N.; Visualization: A.T.N., B.N.

## Data Availability Statement

All code, input data, configuration files, and figures are available at: https://github.com/AlexTaiNguyen006/scfa-dual-model-paper. The pipeline is fully executable end-to-end via a single Makefile using the provided conda environment specification.

